# One prophage WO gene rescues cytoplasmic incompatibility in *Drosophila melanogaster*

**DOI:** 10.1101/300269

**Authors:** J. Dylan Shropshire, Jungmin On, Emily M. Layton, Helen Zhou, Seth R. Bordenstein

**Author notes:** Correspondence to: J. Dylan Shropshire, Nashville, TN, 37235; Seth R. Bordenstein, Nashville, TN, 37235.

## Abstract

*Wolbachia* are maternally-inherited, intracellular bacteria at the forefront of vector control efforts to curb arbovirus transmission. In international field trials, the cytoplasmic incompatibility (CI) drive system of *w*Mel *Wolbachia* is deployed to replace target vector populations, whereby a *Wolbachia*– induced modification of the sperm genome kills embryos. However, *Wolbachia* in the embryo rescue the sperm genome impairment, and therefore CI results in a strong fitness advantage for infected females that transmit the bacteria to offspring. The two genes responsible for the *w*Mel-induced sperm modification of CI, *cifA* and *cifB*, were recently identified in the eukaryotic association module of prophage WO, but the genetic basis of rescue is unresolved. Here we use transgenic and cytological approaches to demonstrate that *cifA* independently rescues CI and nullifies embryonic death caused by *w*Mel *Wolbachia* in *Drosophila melanogaster*. Discovery of *cifA* as the rescue gene and previously one of two CI induction genes establishes a new ‘Two-by-One’ model that underpins the genetic basis of CI. Results highlight the central role of prophage WO in shaping *Wolbachia* phenotypes that are significant to arthropod evolution and vector control.

**Significance Statement:** The World Health Organization recommended pilot deployment of *Wolbachia-*infected mosquitoes to curb viral transmission to humans. Releases of mosquitoes are underway worldwide because *Wolbachia* can block replication of these pathogenic viruses and deterministically spread by a drive system termed cytoplasmic incompatibility (CI). Despite extensive research, the underlying genetic basis of CI remains only half-solved. We recently reported that two prophage WO genes recapitulate the modification component of CI in a released strain for vector control. Here we show that one of these genes underpins rescue of CI. Together, our results reveal the complete genetic basis of this selfish trait and pave the way for future studies exploring WO prophage genes as adjuncts or alternatives to current control efforts.

## Introduction

*Wolbachia* are an archetype of maternally-inherited, intracellular bacteria. They occur in an estimated 40-52% of arthropod species (1, 2) and 47% of the Onchocercidae family of filarial nematodes (3), making them the most widespread bacterial symbiont in the animal kingdom (2). In arthropods, *Wolbachia* mainly reside in the cells of the reproductive tissues, transmit transovarially (4), and often commandeer host fertility, sex ratios, and sex determination to enhance their maternal transmission via male-killing, feminization, parthenogenesis, or cytoplasmic incompatibility (CI) (5, 6).

Discovered nearly half a century ago (7), *Wolbachia*-induced CI is the most common reproductive modification and results in embryonic lethality when an infected male mates with an uninfected female, but this lethality is rescued when the female is likewise infected (8). As such, rescue provides a strong fitness advantage to infected females, the transmitting sex of *Wolbachia* (9–11). Alone, CI-induced lethality is deployed in vector control studies to crash the resident uninfected mosquito population through release of *Wolbachia*-infected males (12–17). Together, CI-induced lethality and rescue constitute a microbial drive system that is used in field studies worldwide to stably replace an uninfected mosquito population with an infected one via release of male and females harboring *w*Mel *Wolbachia* (18), which confer resistance against dengue and Zika viruses (19, 20). The efficacy of this drive system for spreading *Wolbachia* in target populations critically depends on *Wolbachia’s* ability to rescue its own lethal modification of the sperm.

While CI is gaining momentum as a natural, sustainable, and inexpensive tool for vector control, the genes that underpin this microbial adaptation are not fully known. Our previous screen of *Wolbachia* genomes and transcriptomes from infected ovaries identified two adjacent genes, *cifA* and *cifB*, from the *w*Mel strain in *Drosophila melanogaster* as the only genes consistently associated with CI (21). These two genes occur in the eukaryotic association module of prophage WO (22), and they together recapitulate CI when dually expressed in uninfected male flies (21, 23). Each gene alone is incapable of inducing CI (21), and the rescue gene remains unknown. As *cifA* and *cifB* are the only two *w*Mel genes associated with CI, we previously hypothesized that the CI induction and rescue genes might be the same (21). Here we test the hypothesis that transgenic expression of *cifA* and/or *cifB* genes from *w*Mel *Wolbachia* in ovaries can rescue CI and nullify the associated embryonic defects in *D. melanogaster*.

## Results and Discussion

Since *Wolbachia* cannot be genetically transformed, we first tested the ability of *cifA* to transgenically rescue wild type CI using a GAL4-UAS system for tissue-specific expression in uninfected *D. melanogaster* females. As such, we conducted the transgenic experiments under the control of either *nos-*GAL4-*tubulin* in uninfected germline stem cells or maternal triple driver, MTD-GAL4, to drive higher transgene expression throughout oogenesis. In transcriptomes of *w*Mel-infected *D. melanogaster*, *cifA* is a highly expressed prophage WO gene (24). MTD-GAL4 utilizes two *nos*-GAL4 driver variants (including *nos*-GAL4-*tubulin*) and an ovarian tumor driver (25). Control CI and rescue crosses with either driver yielded the expected hatching rates. Crosses between infected males and uninfected females expressing *cifA* under the control of MTD-GAL4 showed a markedly significant increase in embryonic hatching relative to *cifA* expression under *nos*-GAL4-*tubulin* and at levels similar to that in control rescue crosses (Fig. 1A). These results are consistent with complete rescue of CI by *cifA*, in association with increased expression throughout the developing egg chambers. Similar results with *nos*-GAL4-*tubulin* expression in uninfected ovarian germline stem cells resulted in a small increase in hatch rate that was inconsistently significant among replicates (Fig. S1). An analysis of *cifA* gene expression reveals MTD-GAL4 associates with a three-order-of-magnitude increase over *nos*-GAL4-*tubulin*, supporting strength of expression as a factor for rescue (Fig. 1B).

**Fig. 1.**
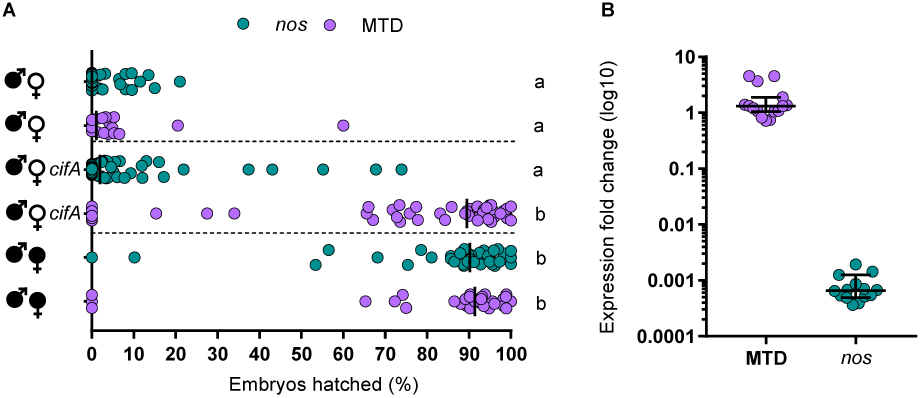
*cifA* rescues cytoplasmic incompatibility when it is highly expressed throughout oogenesis. (A) Hatch rate assays were conducted with transgenic expression of *cifA* under the control of *nos*-GAL4-*tubulin* or MTD-GAL4 drivers. Each dot represents a replicate. Rescue occurred only under MTD-GAL4 expression. Horizontal dotted lines from top to bottom separate cross types with CI, *cifA* expression, and rescue. *Wolbachia* infections are represented by filled sex symbols and expressed genes are noted to the right of the corresponding sex. n=27-59 for each experimental cross across two experiments (both shown). Vertical bars represent medians, and letters to the right indicate significant differences based on α=0.05 calculated by Kruskal-Wallis and Dunn’s test for multiple comparisons. (B) Expression fold change of *cifA* relative to the *Drosophila* housekeeping gene Rp49 was determined on a subset of abdomens from female expressing *cifA* via MTD-GAL4 or *nos*-GAL4-*tubulin* with 2^-∆∆Ct^. Horizontal bars represent medians with 95% confidence intervals, and letters above indicate significance based on a Mann-Whitney test. In both cases, statistical comparisons are between all groups. Exact p-values are provided in Table S2. Hatch rate experiments testing expression of *cifA* under MTD-GAL4 or *nos*-GAL4-*tubulin* have been repeated four and five times respectively.

We expanded our evaluation of *cif* gene expression under the control of MTD-GAL4 in uninfected females to test if *cifB* alone or in combination with *cifA* impacts CI penetrance. As expected, infected males crossed to either uninfected females or females transgenically expressing *cifB* under MTD-GAL4 yielded similar CI penetrance (Fig. 2). These results suggest that *cifB* does not rescue CI when transgenically expressed in the ovaries, and its CI-related function is specific to testes. In contrast, MTD-GAL4 expression of *cifA*, by itself or in combination with *cifB*, significantly rescued CI to levels comparable to rescue by infected females (Fig. 2). These results are consistent with *cifA* independently functioning as the rescue factor and suggest that *cifB* does not inhibit *cifA*’s ability to rescue CI. As *Wolbachia* can induce phenotypes known to bias sex ratios, we collected the surviving offspring from the transgenic and control rescue crosses and sexed them to demonstrate normal sex ratios, indicating that rescue was not sex-specific (Fig. S2).

**Fig. 2.**
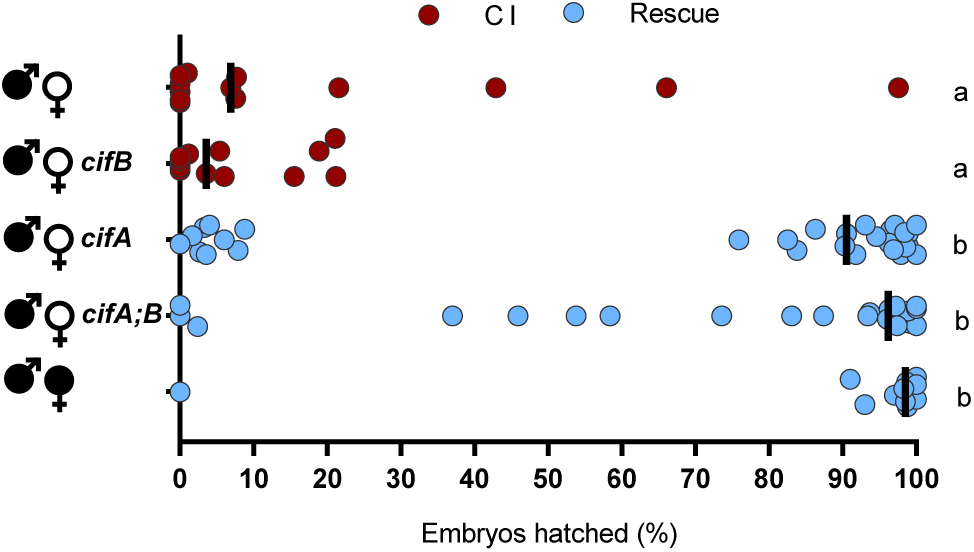
Rescue of cytoplasmic incompatibility is specific to *cifA*. Hatch rate assays were conducted with transgenic expression of *cifA*, *cifB*, and *cifA;B* using the MTD-GAL4 driver for expression throughout oogenesis. Each dot represents a replicate. *Wolbachia* infections are represented by filled sex symbols and expressed genes are noted to the right of the corresponding sex. n=11-29 for each experimental cross. Vertical bars represent medians, and letters to the right indicate significant differences based on α=0.05 calculated by Kruskal-Wallis and Dunn’s test for multiple comparisons. Statistical comparisons are between all groups. Exact p-values are provided in Table S2. Hatch rate experiments testing expression of *cifA* under MTD-GAL4 have been repeated four times.

Next, we tested if the canonical cytological defects observed in early CI embryos (early mitotic failure, chromatin bridging, and regional mitotic failure (26)) were nullified under *cifA*-induced rescue. We examined embryos from control and transgenic crosses after 1-2 h of development and binned their cytology into one of five phenotypes as previously established for *D. melanogaster* CI (21). Nearly half of CI-induced lethality in embryos is the result of embryonic arrest during advanced developmental stages in Dipteran species (27–30). As expected, the control CI cross yielded high levels of all three CI-associated defects, and the embryos from the control rescue cross developed with significantly fewer abnormalities (Fig. 3). MTD-GAL4 transgene expression of *cifA* in uninfected females, either alone or dually expressed with *cifB*, resulted in significantly fewer cytological defects (Fig. 3). These effects were not seen with transgene *cifB* expression, again validating that *cifA* alone can recapitulate wild type rescue by *Wolbachia*.

**Fig. 3.**
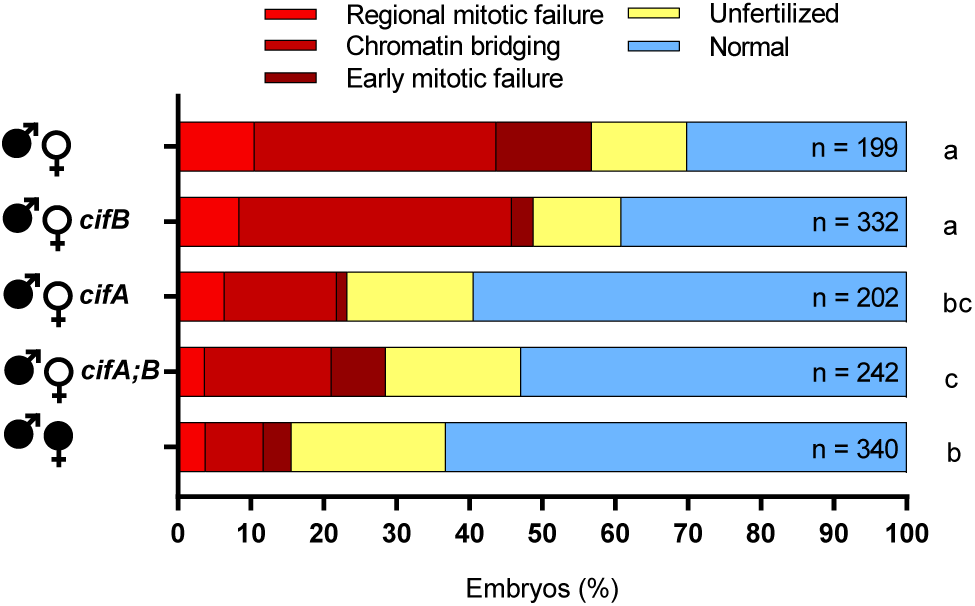
*cifA* rescues embryonic defects caused by cytoplasmic incompatibility. The number of embryos with each cytological phenotype resulting from the indicated crosses is shown. All replicate crosses were conducted in parallel and with sisters from the experiment in Fig 2. *cifA*, *cifB*, and *cifA;B* transgene expression was under the control of MTD-GAL4. *Wolbachia* infections are represented by filled sex symbols and expressed genes are noted to the right of the corresponding sex. Letters to the right indicate significant differences based on α=0.05 calculated by pair-wise chi-square analyses comparing defects (all shades of red) against normal (blue) with Bonferroni adjusted p-values. Exact p-values are provided in Table S2. This experiment has been conducted once.

These data are in contrast with previous work reporting the inability to transgenically rescue CI in *D. melanogaster (23)*; however, there are three critical differences between the studies. First, *w*Pip’s homologs from *Culex pipiens* were used in the prior work instead of *w*Mel’s *cif* genes from *D. melanogaster* here. Thus, differences in host background interactions could explain the discrepancy. Second, a T2A sequence for the wPip gene homologs was used to allow for bicistronic expression, but ribosome skipping results in a C-terminal sequence extension to the first protein and a proline addition to the second protein that generates sequence artifacts and could alter function (31). Finally, different insertion sites are capable of different levels of expression due to their local chromatin environment (32), thus the chosen sites may produce insufficient product to cause rescue, as was the case in our study when *cifA* was driven by *nos*-GAL4-*tubulin*.

*cifA* encodes a putative catalase-rel function, sterile-like transcription factor (STE) domains, and a domain of unknown function (DUF3243) that shares homology with a putative Puf-family RNA binding domain in *cifA*-like homologs (33), whereas *cifB* has nuclease and deubiquitilase domains (23, 33). Only the deubiquitilase annotation has been functionally tested and confirmed(23). Based on subcellular localization (PSORTb) and transmembrane helix predictors (TMbase), CifA is a cytoplasmic protein without transmembrane helices (Fig. S3). Codon-based and Fisher’s exact tests of neutrality demonstrate that closely-related (76.2-99.8% pairwise nucleotide identity) Type I CifA homologs (21) largely evolve by purifying selection (Fig. S4a, b), and sliding window analyses (SWAKK and JCoDA) reveal that purifying selection is strongest on the catalase-rel domain and the unannotated region at the N-terminus, with considerably weaker purifying selection on the putative DUF3243 and STE domains (Fig. 4; Fig. S4c). This is supported by prior work reporting stronger amino acid conservation within the Type I CifA N-terminus relative to the C-terminus (33).

**Fig. 4.**
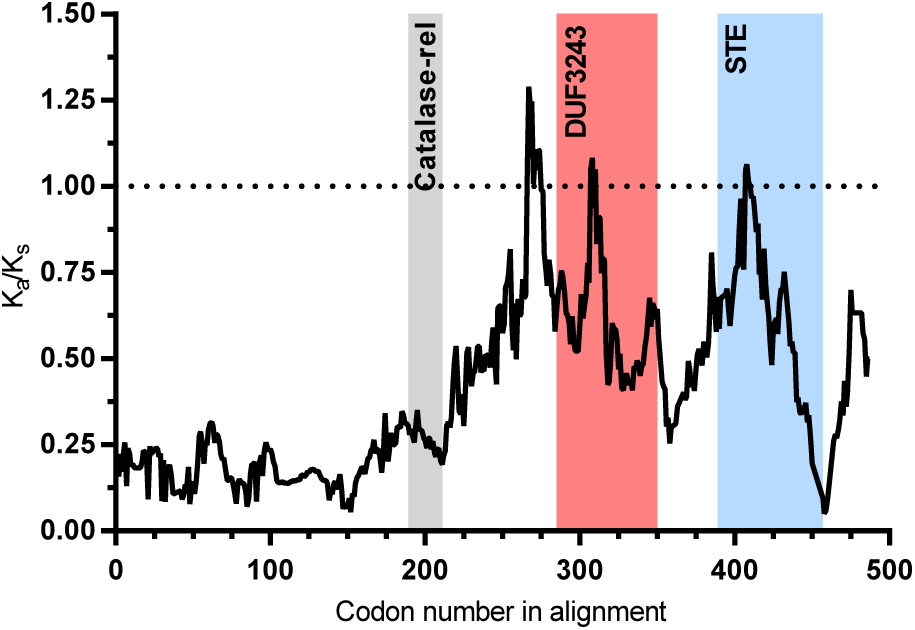
Ka/Ks sliding window analysis identifies *cifA* regions evolving under negative selection. A sliding window analysis of Ka/Ks ratios between *cifA* homologs from *w*Mel and *w*Ha rejects the neutral expectation of Ka/Ks = 1 using a 25 amino acid sliding window across most of *cifA*. Strong purifying selection is observed in several *cifA* regions including the sequence preceding the Catalase-rel domain. Shaded regions denote previously described protein domain predictions (33).

These findings illustrate that the *Wolbachia* prophage WO gene *cifA* recapitulates rescue of wild type CI. As *cifA* is one of two genes involved in induction of CI, results support the hypothesis that a gene involved in CI induction is also the rescue gene (21). In addition, transgenic expression of *cifA* in yeast inhibits a temperature-dependent growth defect caused by *cifB* expression (23). The discovery that CI is induced by *cifA* and *cifB* and rescued by *cifA* motivates a new modification-rescue model of CI where two genes act as the CI modification factors (in the male), and one of these same genes acts as the rescue factor (in the female). This ‘Two-by-One’ model posits that each strain of *Wolbachia* has its own set of *cifA*- and *cifB*-associated CI modifications and one *cifA* rescue factor. The different roles of *cifA* in CI and rescue is intriguing. We predict that the function of *cifA* is dependent on differential tissue localization of gene products in male and female reproductive systems and/or alternate post-translational modification in testes/sperm (CI) versus in ovaries/embryoes (rescue). Moreover, one could speculate that the putative antioxidant catalase-rel domain of the CifA protein acts as a functional switch in the presence of reactive oxygen species, known to be higher in *Wolbachia*-infected testes (34), whereas the Puf-family RNA binding domain and STE are involved in RNA binding and transcriptional (mis)regulation of an unknown host factor.

It has been hypothesized that divergence in modification and rescue genes leads to bidirectional CI (21, 37, 38), which is a reciprocal incompatibility between males and females infected with different *Wolbachia* strains (7, 39–42). Comparative genomic analyses of *cifA* and *cifB* genes reveal extremely high levels of amino acid divergence (21), strong codivergence (21, 33), and recombination (38), consistent with the very rapid evolution of bidirectional CI across *Wolbachia* that can contribute to reproductive isolation and speciation (42, 43). Indeed, divergence of the *cifA* and *cifB* genes into several phylogenetic types correlates with bidirectional CI patterns in *Drosophila* and *Culex* (21, 38). There are at least two explanations for how simple genetic changes in these genes can contribute to bidirectional CI. First, a single mutation in the *cifA* gene could produce variation in the modification and rescue components that render two *Wolbachia* strains incompatible. For instance, given an ancestral and derived allele of *cifA*, males and females with *Wolbachia* carrying the same *cifA* allele are compatible; however, males with *Wolbachia* carrying the ancestral *cifA* allele cause a sperm modification that is unable to be rescued by embryos with *Wolbachia* carrying the derived *cifA* allele, and vice versa. Thus, a single mutation in *cifA* alone can enable the switch from being compatible to incompatible *Wolbachia*. Second, mutations in both *cifA* and *cifB* are required for the evolution of bidirectional CI. For example, CifA-CifB protein binding (23) and/or differential localization in the sperm and egg may underpin bidirectional CI between *Wolbachia* strains. In this model, amino acid divergence in the Cif proteins may contribute to weakened binding, which in turn yields *Wolbachia* strains incapable of CI but capable of rescuing the ancestral variant (44, 45). A compensatory substitution in the other Cif protein could in theory restore binding and yield bidirectional incompatibility with the ancestral Cif variants. Codivergence between amino acid sequences of these proteins is consistent with this model. Under both models, the presence of multiple WO prophages carrying *cifA* genes may also promote incompatibilities through the production of multiple CI product complexes simultaneously (21). In support of these hypotheses, complex diversification and duplication of *cifA* and *cifB* has been reported in *Drosophila* and *C. pipiens* that harbor a variety of incompatible *Wolbachia* strains (21, 38).

In conclusion, our findings reveal the connected genetic basis of CI and rescue and highlight the fundamental impact of prophage genes on the adaptive phenotypes of an obligate intracellular bacteria. In addition to genetically dissecting this widespread form of reproductive parasitism and microbial drive, we also establish a new, Two-by-One model to explain the modification and rescue components of CI. Finally, beneficial applications of CI and rescue genes as transgenic drive constructs may be possible as adjuncts or alternatives to pest control or vector control strategies currently deploying *Wolbachia-*infected mosquitoes (15–18).

## Materials and Methods

### Fly rearing and strains

*D. melanogaster* stocks *y*^1^*w*^*^ (BDSC 1495), *nos-*GAL4-*tubulin* (BDSC 4442), MTD-GAL4 (containing *nos*-GAL4-*tubulin*, nos-GAL4-VP16, and *otu*-GAL4-VP16; BDSC 31777), and UAS transgenic lines homozygous for *cifA*, *cifB*, and *cifA;B* (21) were maintained at 12:12 light:dark at 25 ° C and 70% relative humidity (RH) on 50 ml of a standard media. GAL4 lines were found to be infected with *w*Mel *Wolbachia*, and uninfected lines were produced through tetracycline treatment as previously described (21). Infection status was frequently confirmed via PCR using WolbF and WolbR3 primers (46). During virgin collections, flies were stored at 18 ° C overnight to slow eclosion rate, and virgin flies were kept at room temperature.

### Hatch rate and sex ratio assays

Virgin MTD-GAL4 females were collected for the first 3 days of emergence and aged 9-11 days before crossing to non-virgin homozygous UAS (*cifA*, *cifB*, or *cifA;B*) males. The start of collections for the maternal and paternal lineages were staggered by 7 days. Single pair matings occurred in an 8 oz bottle, and a grape-juice agar plate was smeared with yeast and affixed to the opening with tape. The flies and bottles were then stored at 25 ° C and 70% RH for 24 h at which time the plates were replaced with freshly smeared plates and again stored for 24 h. Plates were then removed and the number of embryos on each plate were counted and stored. After 30 h the remaining unhatched embryos were counted (Extended Data Fig. 6). The hatch rate was calculated by dividing the number of hatched embryos by the initial embryo count and multiplying by 100. Hatch rate was plotted against clutch size for all rescue crosses conducted in this study to reveal a significant correlation (Fig. S5), and a threshold clutch size for analysis was set equal to exclusion of 99% of plates with a hatch rate of 0 for each genotype (31 for *nos*-GAL4-*tubulin* and 48 for MTD-GAL4). Larvae were moved into vials of standard media and the offspring sex ratio determined after 15-18 days (Fig. S6). Hatch rates testing MTD-GAL4 or *nos-*GAL4-*tubulin* expression of *cifA* were conducted three and four times respectively. Sex ratio experiments were conducted once.

### Gene expression

To compare the level of UAS-*cifA* expression between MTD-GAL4 and *nos*-GAL4-*tubulin* flies, mothers from hatch rate assays were collected after the allotted laying period, abdomens were immediately dissected, and samples were frozen in liquid nitrogen and stored at −80C until processing. RNA was extracted using the Direct-zol RNA MiniPrep Kit (Zymo), DNase treated with DNA-free (Ambion, Life Technologies), and cDNA was generated with SuperScript VILO (Invitrogen). Quantitative PCR was performed on a Bio-Rad CFX-96 Real-Time System using iTaq Universal SYBR Green Supermix (Bio-Rad). Forty cycles of PCR were performed against positive controls (extracted DNA), negative controls (water), RNA, and cDNA with the following conditions: 50 °C 10 min, 95 °C 5 min, 40× (95 °C 10 s, 55 °C 30 s), 95 °C 30 s. Primers used were *cifA* opt and Rp49 forward and reverse (Table S1). Fold expression of UAS-*cifA* relative to the *D. melanogaster* house-keeping gene Rp49 was determined with 2^-∆∆Ct^. This experiment and corresponding hatch rate were performed once.

### Embryo cytology

Flies were collected as described for the hatch rate assays, but with 60 females and 12 males in each bottle with a grape-juice agar plate attached. All flies used were siblings of those from the hatch rate, grape-juice plates replaced as described above, and embryos collected in parallel to egg-laying by hatch rate females. Embryos were collected, dechorionated, washed, methanol fixed, stained with propidium iodide, imaged, and categorized as previously described (21) (Fig. S6). This experiment was performed once.

### Putative cifA localization

The PSORTb v3.0.2 web server (47) was used to predict subcellular localization of the *w*Mel CifA protein to either the cytoplasm, cytoplasmic membrane, periplasm, outer membrane, or extracellular space. A localization score is provided for each location with scores of 7.5 or greater considered probable localizations. The TMpred web server (48) was used to predict transmembrane helices in *w*Mel CifA. TMpred scores were generated for transmembrane helices spanning from inside-to-outside (i-o) and outside-to-inside (o-i), and scores above 500 are considered significant.

### cifA selection analyses

Selection analyses were conducted using four independent tests of selection: codon-based Z-test of neutrality (49), Fisher’s exact test of neutrality (49), Sliding Window Analysis of Ka and Ks (SWAKK) (50), and Java Codon Delimited Alignment (JCoDA) (51). The first two analyses were conducted using the MEGA7 desktop app with a MUSCLE translation alignment generated in Geneious v5.5.9. The SWAKK 2.1 web server and the JCoDA v1.4 desktop app were used to analyze divergence between *w*Mel and *w*Ha *cifA* with a sliding window of 25 or 50 codons and a jump size of 1 codon for SWAKK and 5 codons for JCoDA.

### Statistical analyses

All statistical analyses were conducted in GraphPad Prism (Prism 7 or online tools). Hatch rate and sex ratio statistical comparisons were made using Kruskal-Wallis followed by a Dunn’s multiple comparison test. Expression was compared using a Mann-Whitney test. Correlations between hatch rate and clutch size were determined using Spearman rho. Pair-wise chi-square analyses were used for cytology studies to compare defective and normal embryos followed by generation of Bonferroni adjusted p-values. An unpaired t-test was used for statistical comparison of RNA fold expression. All p-values are reported in Table S2.

### Data availability

All source data and replicate data are available as Supplementary Information along with this publication.

## Acknowledgments

This work was supported by National Institutes of Health (NIH) awards R01 AI132581 and R21 HD086833 to S.R.B., National Science Foundation award IOS 1456778 to S.R.B., and a National Science Foundation Graduate Research Fellowship to J.D.S. Any opinion, findings, and conclusions or recommendations expressed in this material are those of the authors(s) and do not necessarily reflect the views of the National Institutes of Health or the National Science Foundation. Imaging was performed in part using the Vanderbilt University Medical Center Cell Imaging Shared Resources (supported by NIH grants CA68485, DK20593, DK58404, DK59637 and EY08126). We thank Daniel LePage for preliminary work, Sarah R. Bordenstein and Jessamyn Perlmutter for manuscript preparation, and Jared Nordman for kindly providing the MTD-GAL4 line.

## Figure Legends

**Fig. S1.**
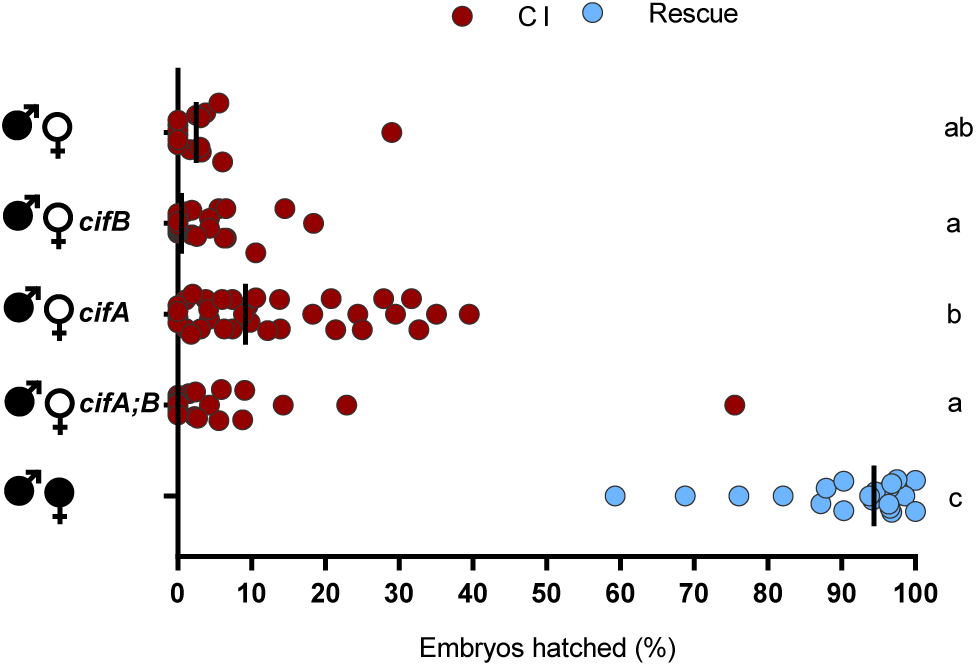
*cifA* transgene expression in germline stem cells fails to elicit rescue. Transgene expression of *cifA*, *cifB*, and *cifA;B* using the *nos*-GAL4-*tubulin* driver does not lead to rescue of cytoplasmic incompatibility. Each dot represents a replicate. *Wolbachia* infections are represented by filled sex symbols, and expressed genes are noted to the right of the corresponding sex. n=15-34 for each experimental cross. Vertical bars represent medians and letters to the right indicate significant differences based on α=0.05 calculated by Kruskal-Wallis and Dunn’s test for multiple comparisons. Statistical comparisons are between all groups. Exact p-values are provided in Table S2.

**Fig. S2.**
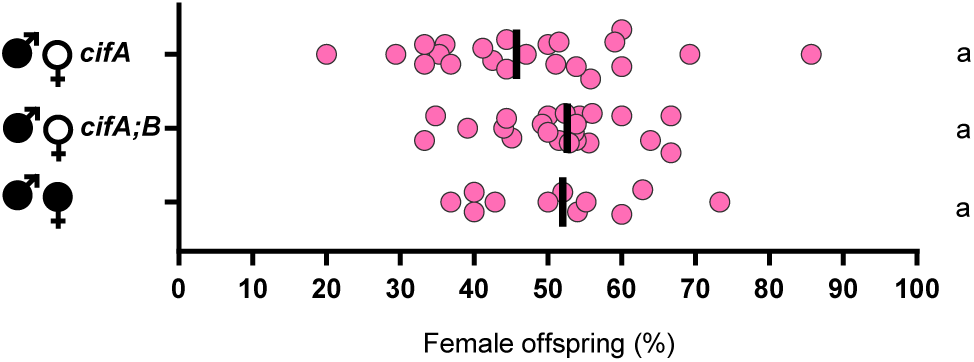
*cifA* does not preferentially rescue one sex over the other. Surviving offspring from the experiment displayed in Figure 2 were collected for adult sex ratio counts. There was no significant difference between any of the crosses. A sex ratio count was not possible for CI crosses due to the low number of surviving offspring. *Wolbachia* infections are represented by filled sex symbols and expressed genes are noted to the right of the corresponding sex. n=11-22 for each experimental cross. Vertical bars represent medians and letters to the right indicate significant differences based on α=0.05 calculated by Kruskal-Wallis and Dunn’s test for multiple comparisons. Statistical comparisons are between all groups. Exact p-values are provided in Table S2. This experiment was conducted once.

**Fig. S3.**
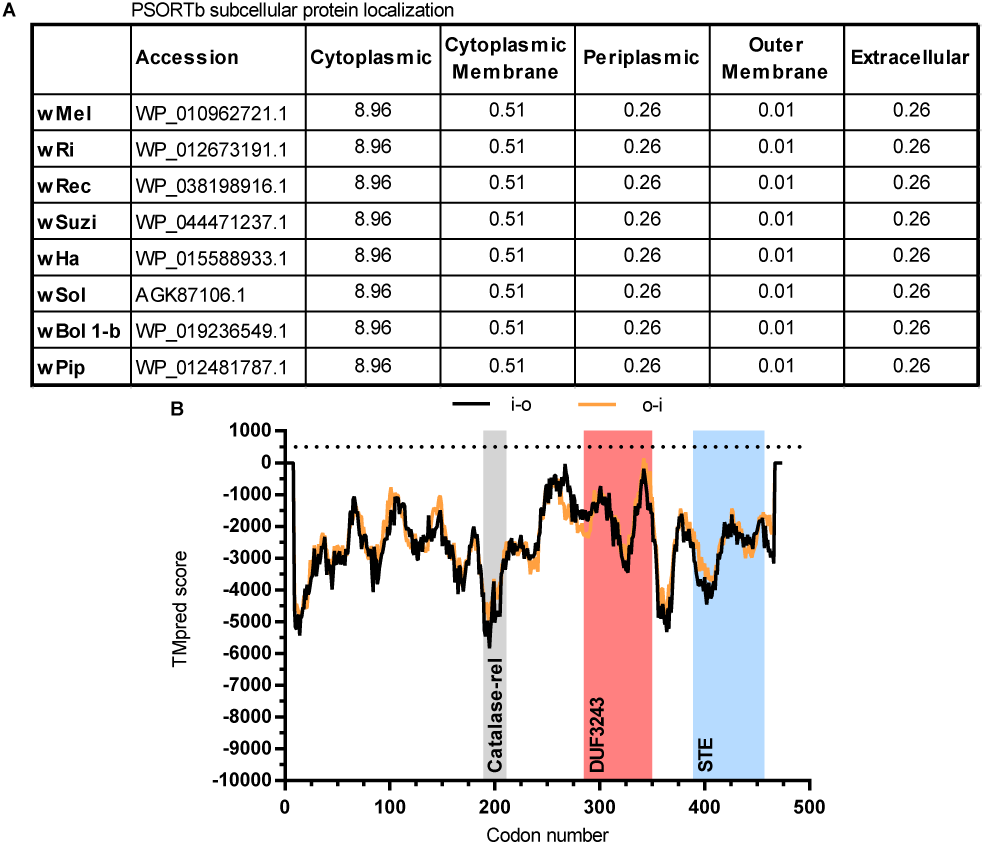
CifA is a putative cytoplasmic protein. (**A**) The PSORTb subcellular protein localization web server was used on Type I CifA proteins to predict the protein’s localization in the *Wolbachia* cell. Predictive scores above 7.5 are accepted to be sufficient to determine a single location of localization and suggest that CifA is a cytoplasmic protein. (**B**) The TMpred web server was used to predict transmembrane helices. TMpred scores exceeding 500 (denoted by horizontal dotted line) are considered significant. TMpred scores were generated for transmembrane helices spanning from inside-to-outside (i-o) and outside-to-inside (o-i). Shaded regions denote previously described protein domain predictions (33).

**Fig. S4.**
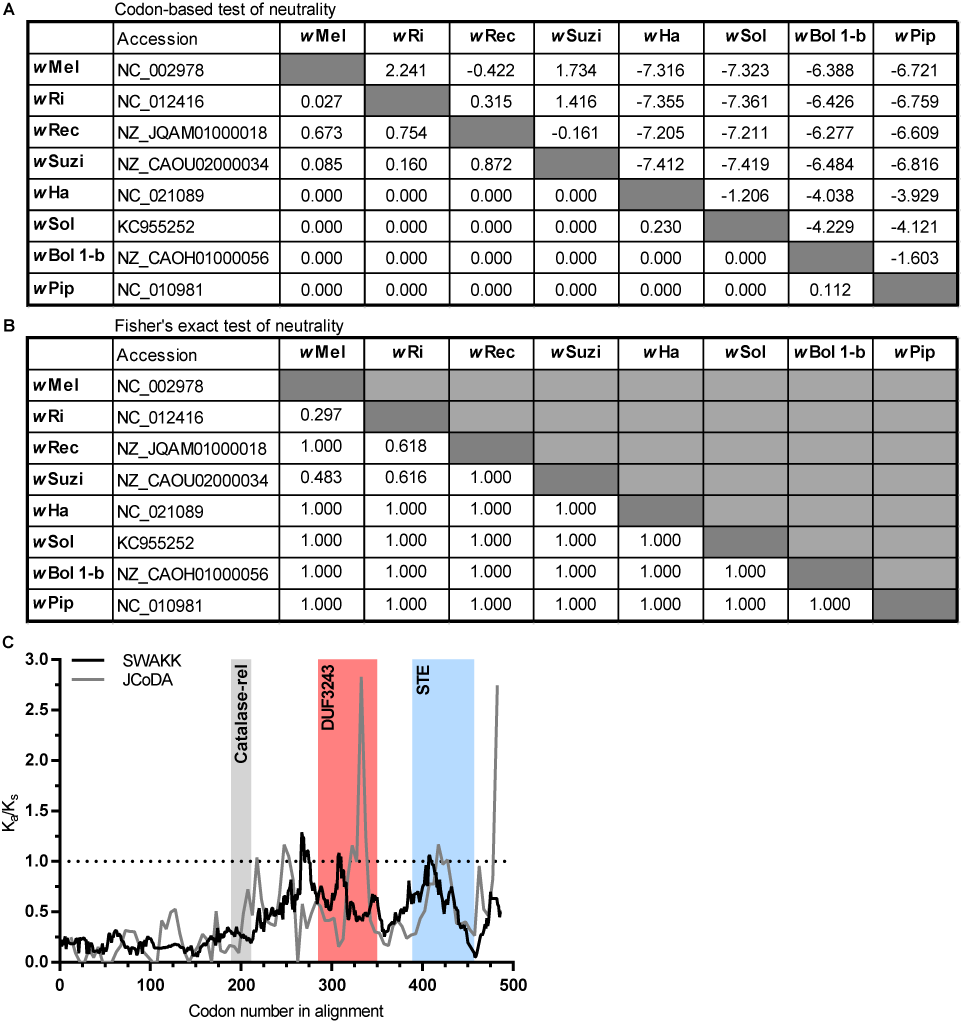
*cifA* regions evolve under negative selection. (**A**) Pairwise codon-based z-tests of selection suggest that regions of the *cifA* gene are not evolving under the neutral expectation of K_a_=K_s_. Values below the diagonal are p-values for where there is a significant departure from neutrality or not. Values above the diagonal are the difference of K_a_-K_s_ in which positive values suggest positive selection and negative values suggest purifying selection. (**B**) Pairwise Fisher’s exact tests of neutrality suggest that *cifA* evolves under purifying selection. Values below the diagonal are p-values. If the p*-*value is less than 0.05, then the null hypothesis of strictly neutral or purifying selection is rejected. If the observed number of synonymous differences per synonymous site exceeds the number of nonsynonymous differences per nonsynonymous site then *MEGA* sets *P* = 1 to indicate purifying selection, rather than positive selection. (**C**) SWAKK and JCoDA were used for sliding window analyses of Ka/Ks ratios between *cifA* homologs of *w*Mel and the bidirectionally incompatible *w*Ha. Both programs were performed with 25 amino acid windows and yield Ka/Ks ratios evident of strong purifying selection in the N-terminus region preceding the Catalase-rel domain and weaker purifying selection beyond it. Shaded regions denote previously described domain predictions (33).

**Fig. S5.**
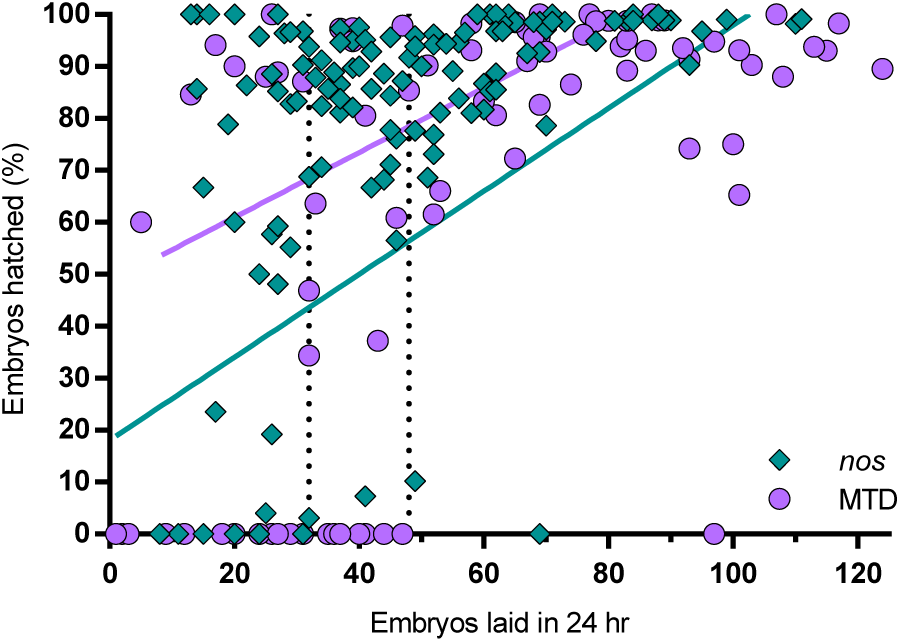
Fertility is related to strain genotype. A meta-analysis of all control rescue crosses (infected male x infected female) without a transgene shows that clutch size and hatch rate are significantly correlated for both the MTD-GAL4 and *nos*-GAL4-*tubulin* genotypes (r = 0.59 and 0.50 for MTD-GAL4 and *nos*-GAL4-*tubulin* respectively), but the two strains have different y-intercepts (4.69 to 31.43 and 39.94 to 59.04 for MTD-GAL4 and *nos*-GAL4-*tubulin* respectively). Each dot represents a replicate where circles and diamonds are MTD-GAL4 (n=91) and *nos-*Gal4*-tubulin* (n=134) respectively. Vertical dotted lines represent embryo counts where 99% of clutch sizes with 0% embryo hatch rate are to the left for *nos*-GAL4-*tubulin* (left line) and MTD-GAL4 (right line). Correlation was assessed with Spearman Rho. A linear regression best-fit line is plotted for each genotype. Exact p-values are provided in Table S2.

**Fig. S6.**
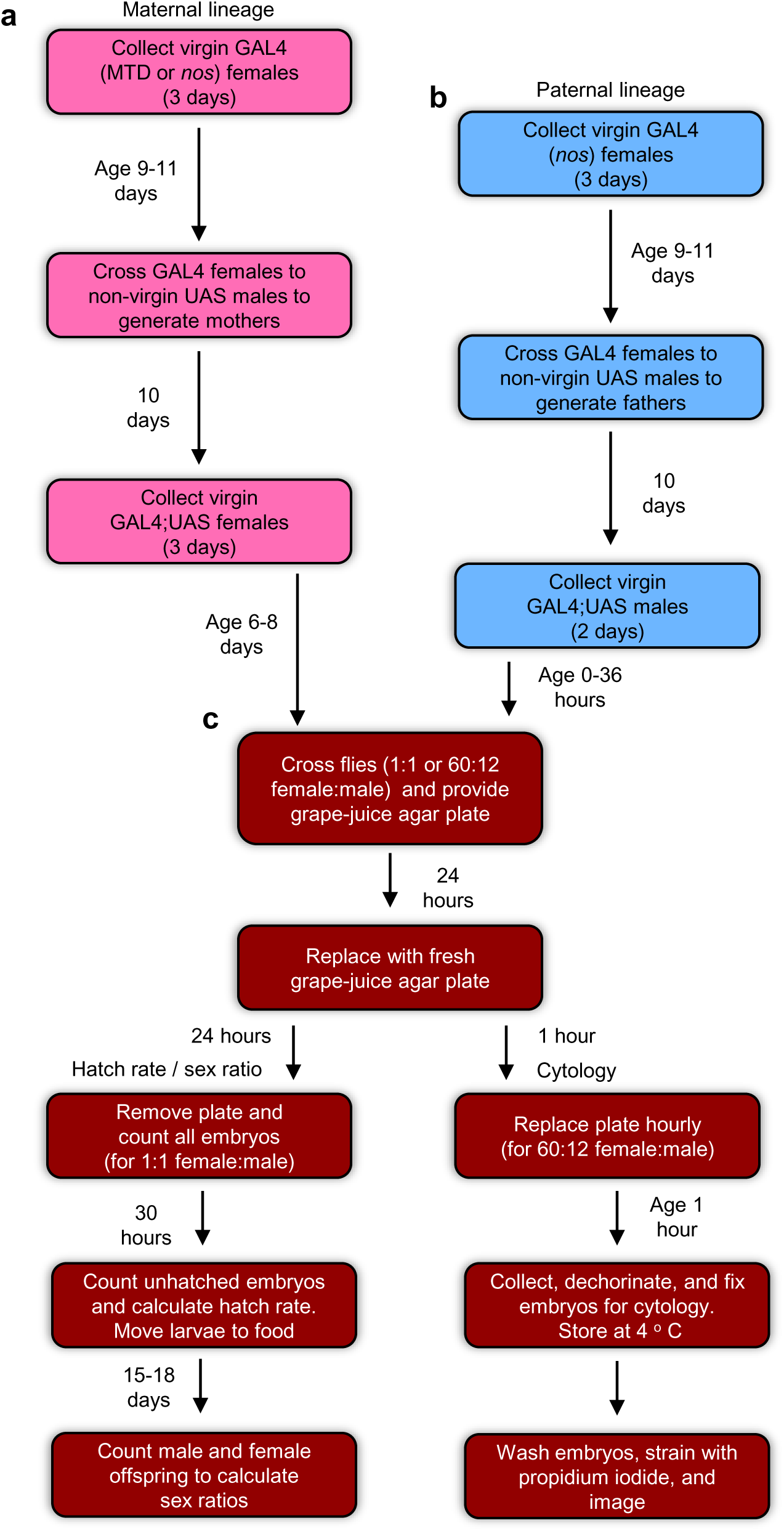
Schematic of experimental methodology. (a) All experimental setups begin with the generation of the maternal lineage (pink), derived from GAL4 driver lines and collected as virgins and aged for 6-8 days till the peak of their fecundity. (b) The paternal lineage (blue) is setup in a stagger such that the males used in the experiment emerge on the day of the experiment. (c) Flies are crossed in a fashion dependent on the ultimate intent, and grape-juice agar plates provided and replaced in a similar manner for all experiments. Sex ratio studies are derived from hatch rate assays.

**Table S1.**
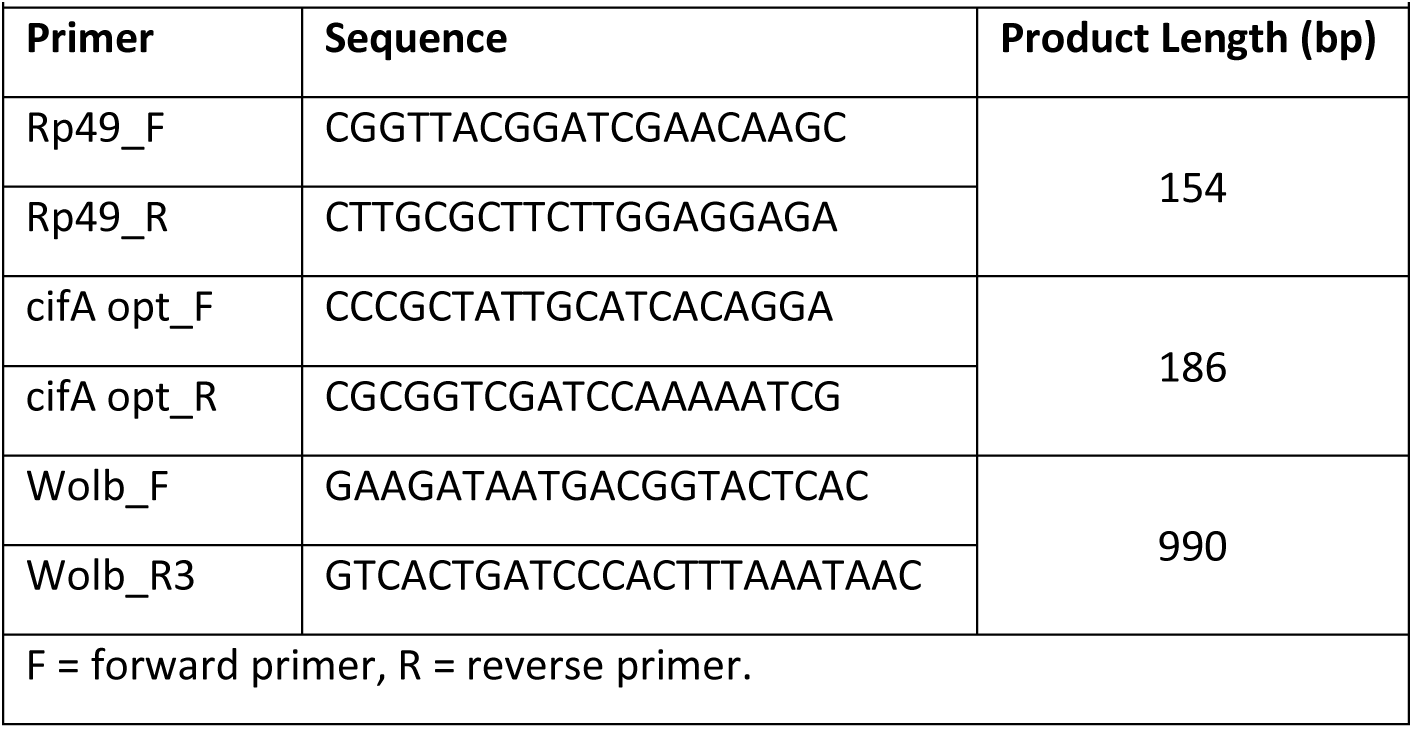
Primers used in this study for RT-qPCR (Fig 1B) or for *Wolbachia* infection checks.

**Table S2.** P-values associated with all statistical comparisons made in main and extended data figures.

|  |  |  |  |
| --- | --- | --- | --- |
| **Figure** | **Comparison** | **p-value** | **Test** |
| Fig. 1A | [M;+]nos;wt x [F;-]MTD;wt vs. [M;+]nos;wt x [F;-]MTD;cifA | **<0.0001** | Kruskal Wallis with Dunn's correction |
|  | [M;+]nos;wt x [F;-]MTD;wt vs. [M;+]nos;wt x [F;+]MTD;wt | **<0.0001** | Kruskal Wallis with Dunn's correction |
|  | [M;+]nos;wt x [F;-]MTD;wt vs. [M;+]nos;wt x [F;-]nos;wt | >0.9999 | Kruskal Wallis with Dunn's correction |
|  | [M;+]nos;wt x [F;-]MTD;wt vs. [M;+]nos;wt x [F;-]nos;cifA | >0.9999 | Kruskal Wallis with Dunn's correction |
|  | [M;+]nos;wt x [F;-]MTD;wt vs. [M;+]nos;wt x [F;+]nos;wt | **<0.0001** | Kruskal Wallis with Dunn's correction |
|  | [M;+]nos;wt x [F;-]MTD;cifA vs. [M;+]nos;wt x [F;+]MTD;wt | >0.9999 | Kruskal Wallis with Dunn's correction |
|  | [M;+]nos;wt x [F;-]MTD;cifA vs. [M;+]nos;wt x [F;-]nos;wt | **<0.0001** | Kruskal Wallis with Dunn's correction |
|  | [M;+]nos;wt x [F;-]MTD;cifA vs. [M;+]nos;wt x [F;-]nos;cifA | **<0.0001** | Kruskal Wallis with Dunn's correction |
|  | [M;+]nos;wt x [F;-]MTD;cifA vs. [M;+]nos;wt x [F;+]nos;wt | >0.9999 | Kruskal Wallis with Dunn's correction |
|  | [M;+]nos;wt x [F;+]MTD;wt vs. [M;+]nos;wt x [F;-]nos;wt | **<0.0001** | Kruskal Wallis with Dunn's correction |
|  | [M;+]nos;wt x [F;+]MTD;wt vs. [M;+]nos;wt x [F;-]nos;cifA | **<0.0001** | Kruskal Wallis with Dunn's correction |
|  | [M;+]nos;wt x [F;+]MTD;wt vs. [M;+]nos;wt x [F;+]nos;wt | >0.9999 | Kruskal Wallis with Dunn's correction |
|  | [M;+]nos;wt x [F;-]nos;wt vs. [M;+]nos;wt x [F;-]nos;cifA | >0.9999 | Kruskal Wallis with Dunn's correction |
|  | [M;+]nos;wt x [F;-]nos;wt vs. [M;+]nos;wt x [F;+]nos;wt | **<0.0001** | Kruskal Wallis with Dunn's correction |
|  | [M;+]nos;wt x [F;-]nos;cifA vs. [M;+]nos;wt x [F;+]nos;wt | **<0.0001** | Kruskal Wallis with Dunn's correction |
| Fig. 1B | nos-GAL4-tubulin vs. MTD-GAL4 cifA expression | **<0.0001** | Mann-Whitney test |
| Fig. 2 | [M;+]nos;wt x [F;-]MTD;wt vs. [M;+]nos;wt x [F;-]MTD;cifA | **0.0379** | Kruskal Wallis with Dunn's correction |
|  | [M;+]nos;wt x [F;-]MTD;wt vs. [M;+]nos;wt x [F;-]MTD;cifB | >0.9999 | Kruskal Wallis with Dunn's correction |
|  | [M;+]nos;wt x [F;-]MTD;wt vs. [M;+]nos;wt x [F;-]MTD;cifA;cifB | **0.0038** | Kruskal Wallis with Dunn's correction |
|  | [M;+]nos;wt x [F;-]MTD;wt vs. [M;+]nos;wt x [F;+]MTD;wt | **0.0006** | Kruskal Wallis with Dunn's correction |
|  | [M;+]nos;wt x [F;-]MTD;cifA vs. [M;+]nos;wt x [F;-]MTD;cifB | **0.0058** | Kruskal Wallis with Dunn's correction |
|  | [M;+]nos;wt x [F;-]MTD;cifA vs. [M;+]nos;wt x [F;-]MTD;cifA;cifB | >0.9999 | Kruskal Wallis with Dunn's correction |
|  | [M;+]nos;wt x [F;-]MTD;cifA vs. [M;+]nos;wt x [F;+]MTD;wt | 0.5436 | Kruskal Wallis with Dunn's correction |
|  | [M;+]nos;wt x [F;-]MTD;cifB vs. [M;+]nos;wt x [F;-]MTD;cifA;cifB | **0.0004** | Kruskal Wallis with Dunn's correction |
|  | [M;+]nos;wt x [F;-]MTD;cifB vs. [M;+]nos;wt x [F;+]MTD;wt | **<0.0001** | Kruskal Wallis with Dunn's correction |
|  | [M;+]nos;wt x [F;-]MTD;cifA;cifB vs. [M;+]nos;wt x [F;+]MTD;wt | >0.9999 | Kruskal Wallis with Dunn's correction |
| Fig. 3 | [M;+]nos;wt x [F;-]MTD;wt vs. [M;+]nos;wt x [F;-]MTD;cifA | **0.0010** | Chi-square with bonferroni adjusted p-value |
|  | [M;+]nos;wt x [F;-]MTD;wt vs. [M;+]nos;wt x [F;-]MTD;cifB | 0.4680 | Chi-square with bonferroni adjusted p-value |
|  | [M;+]nos;wt x [F;-]MTD;wt vs. [M;+]nos;wt x [F;-]MTD;cifA;cifB | **0.0010** | Chi-square with bonferroni adjusted p-value |
|  | [M;+]nos;wt x [F;-]MTD;wt vs. [M;+]nos;wt x [F;+]MTD;wt | **0.0010** | Chi-square with bonferroni adjusted p-value |
|  | [M;+]nos;wt x [F;-]MTD;cifA vs. [M;+]nos;wt x [F;-]MTD;cifB | **0.0010** | Chi-square with bonferroni adjusted p-value |
|  | [M;+]nos;wt x [F;-]MTD;cifA vs. [M;+]nos;wt x [F;-]MTD;cifA;cifB | 1.0000 | Chi-square with bonferroni adjusted p-value |
|  | [M;+]nos;wt x [F;-]MTD;cifA vs. [M;+]nos;wt x [F;+]MTD;wt | 0.5740 | Chi-square with bonferroni adjusted p-value |
|  | [M;+]nos;wt x [F;-]MTD;cifB vs. [M;+]nos;wt x [F;-]MTD;cifA;cifB | **0.0010** | Chi-square with bonferroni adjusted p-value |
|  | [M;+]nos;wt x [F;-]MTD;cifB vs. [M;+]nos;wt x [F;+]MTD;wt | **0.0010** | Chi-square with bonferroni adjusted p-value |
|  | [M;+]nos;wt x [F;-]MTD;cifA;cifB vs. [M;+]nos;wt x [F;+]MTD;wt | **0.0030** | Chi-square with bonferroni adjusted p-value |
| Fig. S1 | [M;+]nos;wt x [F;-]nos;wt vs. [M;+]nos;wt x [F;-]nos;cifA | 0.1534 | Kruskal Wallis with Dunn's correction |
|  | [M;+]nos;wt x [F;-]nos;wt vs. [M;+]nos;wt x [F;-]nos;cifB | >0.9999 | Kruskal Wallis with Dunn's correction |
|  | [M;+]nos;wt x [F;-]nos;wt vs. [M;+]nos;wt x [F;-]nos;cifA;cifB | >0.9999 | Kruskal Wallis with Dunn's correction |
|  | [M;+]nos;wt x [F;-]nos;wt vs. [M;+]nos;wt x [F;+]nos;wt | **<0.0001** | Kruskal Wallis with Dunn's correction |
|  | [M;+]nos;wt x [F;-]nos;cifA vs. [M;+]nos;wt x [F;-]nos;cifB | **0.0204** | Kruskal Wallis with Dunn's correction |
|  | [M;+]nos;wt x [F;-]nos;cifA vs. [M;+]nos;wt x [F;-]nos;cifA;cifB | **0.0306** | Kruskal Wallis with Dunn's correction |
|  | [M;+]nos;wt x [F;-]nos;cifA vs. [M;+]nos;wt x [F;+]nos;wt | **0.0001** | Kruskal Wallis with Dunn's correction |
|  | [M;+]nos;wt x [F;-]nos;cifB vs. [M;+]nos;wt x [F;-]nos;cifA;cifB | >0.9999 | Kruskal Wallis with Dunn's correction |
|  | [M;+]nos;wt x [F;-]nos;cifB vs.<br>[M;+]nos;wt x [F;+]nos;wt | **<0.0001** | Kruskal Wallis with Dunn's correction |
|  | [M;+]nos;wt x [F;-]nos;cifA;cifB vs.<br>[M;+]nos;wt x [F;+]nos;wt | **<0.0001** | Kruskal Wallis with Dunn's correction |
| Fig. S2 | [M;+]nos;wt x [F;-]MTD;cifA vs.<br>[M;+]nos;wt x [F;-]MTD;cifA;cifB | 0.5209 | Kruskal Wallis with Dunn's correction |
|  | [M;+]nos;wt x [F;-]MTD;cifA vs.<br>[M;+]nos;wt x [F;+]MTD;wt | 0.8609 | Kruskal Wallis with Dunn's correction |
|  | [M;+]nos;wt x [F;-]MTD;cifA;cifB<br>vs. [M;+]nos;wt x [F;+]MTD;wt | >0.9999 | Kruskal Wallis with Dunn's correction |
| Fig. S5 | Hatch rate vs clutch size (MTD-GAL4) | **<0.0001** | Spearman's Rho |
|  | Hatch rate vs clutch size (nos-GAL4-tubulin) | **<0.0001** | Spearman's Rho |
| M = male, F = female, + = *Wolbachia* infected, - = *Wolbachia* uninfected, bold p-values = significant |  |  |  |

## References

1. Weinert LA, Araujo-Jnr EV, Ahmed MZ, & Welch JJ (2015) The incidence of bacterial endosymbionts in terrestrial arthropods. Proc Biol Sci 282(1807):20150249.

2. Zug R & Hammerstein P (2012) Still a host of hosts for Wolbachia: analysis of recent data suggests that 40% of terrestrial arthropod species are infected. PLoS One 7(6):e38544.

3. Ferri E, et al. (2011) New insights into the evolution of *Wolbachia* infections in filarial nematodes inferred from a large range of screened species. PLoS One 6(6):e20843.

4. Frydman HM, Li JM, Robson DN, & Wieschaus E (2006) Somatic stem cell niche tropism in *Wolbachia*. Nature 441(7092):509–512.

5. Werren JH, Baldo L, & Clark ME (2008) *Wolbachia*: master manipulators of invertebrate biology. Nat Rev Microbiol 6(10):741–751.

6. LePage D & Bordenstein SR (2013) *Wolbachia*: Can we save lives with a great pandemic? Trends in parasitology 29(8):385–393.

7. Yen J & Barb A (1973) The etiological agent of cytoplasmic incompatability in *Culex pipiens*. Journal of Invertebrate Pathology 22.

8. Serbus LR, Casper-Lindley C, Landmann F, & Sullivan W (2008) The genetics and cell biology of *Wolbachia*-host interactions. Annu Rev Genet 42:683–707.

9. Turelli M & Hoffmann AA (1999) Microbe-induced cytoplasmic incompatibility as a mechanism for introducing transgenes into arthropod populations. Insect Mol Biol 8(2):243–255.

10. Hancock PA, Sinkins SP, & Godfray HC (2011) Strategies for introducing *Wolbachia* to reduce transmission of mosquito-borne diseases. PLoS Negl Trop Dis 5(4):e1024.

11. Hancock PA, Sinkins SP, & Godfray HC (2011) Population dynamic models of the spread of *Wolbachia*. Am Nat 177(3):323–333.

12. Atyame CM, et al. (2011) Cytoplasmic incompatibility as a means of controlling *Culex pipiens* quinquefasciatus mosquito in the islands of the south-western Indian Ocean. PLoS Negl Trop Dis 5(12):e1440.

13. Dobson SL, Bordenstein SR, & Rose RI (2016) *Wolbachia* mosquito control: Regulated. Science 352(6285):526–527.

14. O’Connor L, et al. (2012) Open release of male mosquitoes infected with a *Wolbachia* biopesticide: field performance and infection containment. PLoS Negl Trop Dis 6(11):e1797.

15. Zhang D, Zheng X, Xi Z, Bourtzis K, & Gilles JR (2015) Combining the sterile insect technique with the incompatible insect technique: I-impact of *Wolbachia* infection on the fitness of triple- and double-infected strains of *Aedes albopictus*. PLoS One 10(4):e0121126.

16. Ritchie SA, Townsend M, Paton CJ, Callahan AG, & Hoffmann AA (2015) Application of *w*MelPop *Wolbachia* strain to crash local populations of *Aedes aegypti*. PLoS Negl Trop Dis 9(7):e0003930.

17. Zabalou S, et al. (2004) *Wolbachia*-induced cytoplasmic incompatibility as a means for insect pest population control. Proc Natl Acad Sci U S A 101(42):15042–15045.

18. Hoffmann AA, et al. (2014) Stability of the *w*Mel *Wolbachia* infection following invasion into *Aedes aegypti* populations. PLoS Negl Trop Dis 8(9):e3115.

19. Walker T, et al. (2011) The *w*Mel *Wolbachia* strain blocks dengue and invades caged *Aedes aegypti* populations. Nature 476(7361):450–453.

20. Dutra HL, et al. (2016) *Wolbachia* blocks currently circulating Zika virus isolates in Brazilian *Aedes aegypti* mosquitoes. Cell Host Microbe 19(6):771–774.

21. LePage DP, et al. (2017) Prophage WO genes recapitulate and enhance *Wolbachia*-induced cytoplasmic incompatibility. Nature 543(7644):243–247.

22. Bordenstein SR & Bordenstein SR (2016) Eukaryotic association module in phage WO genomes from *Wolbachia*. Nat Commun 7:13155.

23. Beckmann JF, Ronau JA, & Hochstrasser M (2017) A *Wolbachia* deubiquitylating enzyme induces cytoplasmic incompatibility. Nat Microbiol 2:17007.

24. Gutzwiller F, et al. (2015) Dynamics of *Wolbachia* pipientis Gene Expression Across the *Drosophila melanogaster* Life Cycle. G3 (Bethesda) 5(12):2843–2856.

25. Staller MV, et al. (2013) Depleting gene activities in early *Drosophila* embryos with the “maternal-Gal4-shRNA” system. Genetics 193(1):51–61.

26. Landmann F, Orsi GA, Loppin B, & Sullivan W (2009) *Wolbachia*-mediated cytoplasmic incompatibility is associated with impaired histone deposition in the male pronucleus. PLoS Pathog 5(3):e1000343.

27. Lassy CW & Karr TL (1996) Cytological analysis of fertilization and early embryonic development in incompatible crosses of *Drosophila simulans*. Mechanisms of development 57(1):47–58.

28. Callaini G, Riparbelli MG, Giordano R, & Dallai R (1996) Mitotic defects associated with cytoplasmic incompatibility in *Drosophila simulans*. Journal of Invertebrate Pathology 67(1):55–64.

29. Wright JD & Barr AR (1981) *Wolbachia* and the normal and incompatible eggs of *Aedes polynesiensis* (Diptera: Culicidae). Journal of Invertebrate Pathology 38(3):409–418.

30. Duron O & Weill M (2006) *Wolbachia* infection influences the development of *Culex pipiens* embryo in incompatible crosses. Heredity (Edinb) 96(6):493–500.

31. Donnelly ML, et al. (2001) Analysis of the aphthovirus 2A/2B polyprotein ‘cleavage’ mechanism indicates not a proteolytic reaction, but a novel translational effect: a putative ribosomal ‘skip’. The Journal of general virology 82(Pt 5):1013–1025.

32. Akhtar W, et al. (2013) Chromatin Position Effects Assayed by Thousands of Reporters Integrated in Parallel. Cell 154(4):914–927.

33. Lindsey ARI, et al. (2018) Evolutionary genetics of cytoplasmic incompatibility genes *cifA* and *cifB* in prophage WO of *Wolbachia*. Genome biology and evolution 10(2):435–451.

34. Brennan LJ, Haukedal JA, Earle JC, Keddie B, & Harris HL (2012) Disruption of redox homeostasis leads to oxidative DNA damage in spermatocytes of Wolbachia-infected Drosophila simulans. Insect Mol Biol 21(5):510–520.

35. Penz T, et al. (2012) Comparative genomics suggests an independent origin of cytoplasmic incompatibility in *Cardinium hertigii*. PLoS Genet 8(10):e1003012.

36. Mann E, et al. (2017) Transcriptome Sequencing Reveals Novel Candidate Genes for *Cardinium hertigii*-Caused Cytoplasmic Incompatibility and Host-Cell Interaction. mSystems 2(6).

37. Charlat S, Calmet C, & Mercot H (2001) On the mod resc model and the evolution of Wolbachia compatibility types. Genetics 159(4):1415–1422.

38. Bonneau M, et al. (2018) Culex pipiens crossing type diversity is governed by an amplified and polymorphic operon of Wolbachia. Nature Communications 9(1):319.

39. O’Neill SL & Karr TL (1990) bidirectional incompatability between conspecific populations of drosophila simulans. Nature 348:178–180.

40. Bordenstein SR & Werren JH (2007) Bidirectional incompatibility among divergent *Wolbachia* and incompatibility level differences among closely related *Wolbachia* in *Nasonia*. Heredity 99(3):278–287.

41. Poinsot D, Bourtzis K, Markakis G, Savakis C, & Mercot H (1998) *Wolbachia* transfer from *Drosophila melanogaster* into *D. simulans*: Host effect and cytoplasmic incompatibility relationships. Genetics 150(1):227–237.

42. Bordenstein SR, O’Hara FP, & Werren JH (2001) *Wolbachia*-induced incompatibility precedes other hybrid incompatibilities in *Nasonia*. Nature 409(6821):707–710.

43. Brucker RM & Bordenstein SR (2012) Speciation by symbiosis. Trends in ecology & evolution 27(8):443–451.

44. Zabalou S, et al. (2004) Natural *Wolbachia* infections in the *Drosophila yakuba* species complex do not induce cytoplasmic incompatibility but fully rescue the *w*Ri modification. Genetics 167(2):827–834.

45. Bourtzis K, Dobson SL, Braig HR, & O’Neill SL (1998) Rescuing *Wolbachia* have been overlooked. Nature 391(6670):852–853.

46. Casiraghi M, Anderson TJ, Bandi C, Bazzocchi C, & Genchi C (2001) A phylogenetic analysis of filarial nematodes: comparison with the phylogeny of *Wolbachia* endosymbionts. Parasitology 122 Pt 1:93–103.

47. Yu NY, et al. (2010) PSORTb 3.0: improved protein subcellular localization prediction with refined localization subcategories and predictive capabilities for all prokaryotes. Bioinformatics (Oxford, England) 26(13):1608–1615.

48. Hofmann K (1993) TMbase-A database of membrane spanning proteins segments. Biol. Chem. Hoppe-Seyler 374:166.

49. Kumar S, Stecher G, & Tamura K (2016) MEGA7: Molecular Evolutionary Genetics Analysis Version 7.0 for Bigger Datasets. Molecular biology and evolution 33(7):1870–1874.

50. Liang H, Zhou W, & Landweber LF (2006) SWAKK: a web server for detecting positive selection in proteins using a sliding window substitution rate analysis. Nucleic Acids Res 34(Web Server issue):W382–384.

51. Steinway SN, Dannenfelser R, Laucius CD, Hayes JE, & Nayak S (2010) JCoDA: a tool for detecting evolutionary selection. BMC Bioinformatics 11:284.

